# African *Salmonella* Typhimurium sequence type 313 lineage 2 evades MAIT cell recognition by overexpressing RibB

**DOI:** 10.1101/762955

**Authors:** Lorena Preciado-Llanes, Anna Aulicino, Rocío Canals, Patrick Moynihan, Xiaojun Zhu, Ndaru Jambo, Tonney Nyirenda, Innocent Kadwala, Siân V. Owen, Natacha Veerapen, Gurdyal S. Besra, Melita A. Gordon, Jay C. D. Hinton, Giorgio Napolitani, Mariolina Salio, Alison Simmons

## Abstract

Mucosal-associated invariant T (MAIT) cells are a subset of innate T lymphocytes activated by bacteria that produce vitamin B2 metabolites. Mouse models of infection have demonstrated a role for MAIT cells in antimicrobial defence. However, proposed protective roles of MAIT cells in human infections remain unproven and clinical conditions associated with a selective absence of MAIT cells have not been identified. We report that typhoidal and non-typhoidal *S. enterica* strains generally activate MAIT cells. However, African invasive disease-associated multidrug-resistant *S.* Typhimurium sequence type 313 lineage 2 strains escape MAIT cell recognition through overexpression of *ribB*, a bacterial gene that encodes the 4-dihydroxy-2-butanone 4-phosphate synthase enzyme of the riboflavin biosynthetic pathway. This MAIT cell-specific phenotype did not extend to other innate lymphocytes. We propose that *ribB* overexpression is an evolved trait that facilitates evasion from immune recognition by MAIT cells and contributes to the invasive pathogenesis of *S.* Typhimurium sequence type 313 lineage 2 *in vivo*.

## INTRODUCTION

The Gram-negative bacterium *Salmonella enterica spp*. comprises many serovars which are closely related phylogenetically but cause very different disease presentations and distinct immune responses in immunocompetent hosts [1], [2]. Infection by the human restricted *Salmonella* typhoidal serovars (*S.* Typhi and *S.* Paratyphi), results in a severe systemic disease called enteric fever. In contrast, nontyphoidal serovars originating from zoonotic reservoirs such as *S.* Typhimurium and *S.* Enteritidis, cause self-limiting diarrhoeal disease in healthy individuals [1]–[3]. Multi-drug resistant *S.* Typhimurium strains of a distinct multilocus sequence type 313 recently emerged in sub-Saharan Africa. *S.* Typhimurium sequence type 313 is associated with invasive blood stream infections in immunocompromised individuals and is distinct from the S. Typhimurium strains that cause gastroenteritis globally.

Since it was first reported in 2009 [4], the *S.* Typhimurium sequence type 313 clade has become the major cause of invasive nontyphoidal *Salmonella* (iNTS) disease in Africa [5], [6], and comprises two sub-clade lineages [6], termed lineages 1 and 2. Bacteraemia by iNTS causes an estimated 681,000 deaths annually worldwide [7], primarily in Africa, among young children with recent malaria, malarial anaemia or malnutrition and in adults afflicted with HIV, among whom recurrent disease is also common [1], [5], [8]–[11]. *S.* Typhimurium sequence type 313 isolates have rarely been reported outside of Africa [4] and African sequence type 313 blood isolates are genetically distinct from rare diarrhoeal sequence type 313 isolates found in the United Kingdom [12] or Brazil [13]. Genotypic and phenotypic analyses of several clinical isolates of the two well-described sequence type 313 lineages identified signatures of metabolic adaptation and unique enteropathogenesis in animal models, consistent with adaptation to invasive disease in an immunocompromised human population [4], [14], [15].

B and T cell responses can mediate a protective role in mouse models of *Salmonella* infection. B cells provide the first line of defence at mucosal sites to restrain systemic dissemination, while T cells are needed for *Salmonella* clearance [16]–[18]. Cross-reactive and serovar-specific MHC restricted T cell responses have been well characterised in humans [19]–[24]. *Salmonella* can also induce activation of non-MHC restricted T cells, specifically γδ T cells, invariant Natural Killer T cells (iNKT) and Mucosal-associated invariant T (MAIT) cells [25]–[27], although their protective role remains undetermined.

MR1-restricted MAIT cells comprise a highly conserved class of semi invariant T cells, bridging innate and adaptive immunity [28]. The MHC class I-like molecule MR1, bound to derivatives of vitamin B2 intermediates, activates MAIT cells [29]. This process can drive antibacterial activity, *in vitro* and *in vivo*, and correlates with the presence of the vitamin B2 biosynthetic pathway in several commensal and pathogenic bacteria and fungal species (reviewed in [30]). Similar to iNKT and γδ T cells, MAIT can be activated by cytokines (IL-12, IL-18, type I IFN) independently of their TCR engagement [31]. The ability of MAIT cells to recognise *S.* Typhimurium-infected targets [32] prompted the identification, within bacterial supernatants, of the potent MAIT cell agonists (lumazine and pyrimidines), derivatives of the vitamin B2 intermediate 5-A-RU [29], [33]. Following intranasal infection with *S.* Typhimurium, murine MAIT cells become activated and accumulate in the lungs [26]. Human challenge studies with typhoidal serovars (*S.* Typhi and *S.* Paratyphi A) also demonstrated sustained MAIT cell activation and proliferation at peak of infection [34], [35].

While many commensal and pathogenic bacteria possess the riboflavin biosynthetic pathway, the levels of resulting MAIT stimulation varies [36], [37], perhaps reflecting the influence of microenvironment on bacterial metabolism and antigen availability. The ability of MAIT cells to recognise and respond to several isolates of the same pathogen may also vary depending on metabolic differences between isolates [38].

We hypothesized that MAIT cells contributed to the cellular response to *Salmonella enterica* serovars responsible for invasive disease, and examined the ability of MAIT cells to recognise and respond to different *S. enterica* serovars associated to invasive disease in Africa. Here, we demonstrate that *S.* Typhimurium sequence type 313 lineage 2 isolates escape MAIT cell recognition through overexpression of RibB, a bacterial enzyme of the riboflavin biosynthetic pathway. Our results suggest that MAIT cell immune protection represents an important ‘evolutionary bottleneck’ for the pathogen.

## RESULTS

### Identification of cellular responses to multiple *Salmonella enterica* subsp *enterica* serovars

To identify potential differences in the response of innate and adaptive T cells to distinct *Salmonella* pathovars, we focused on two pathovariants of *S.* Typhimurium that are responsible for different types of human disease. *S.* Typhimurium sequence type 313 is associated with invasive disease among immunocompromised individuals in Africa and the representative isolate is D23580 (STM-D23580). *S.* Typhimurium sequence type 19 causes non-invasive diarrhoeal infections in immunocompetent individuals globally (representative isolate is LT2, designated STM-LT2). Peripheral blood mononuclear cells (PBMC) isolated from healthy donors were infected with both *S.* Typhimurium pathovariants, and *S.* Typhi strain Ty2 (ST-Ty2) was used to represent a more distantly-related serovar that causes invasive disease in immunocompetent individuals. *Escherichia coli (E. coli)* was included as unrelated control. Upon infection, PBMC were incubated overnight in the presence of brefeldin A to permit intracellular cytokine accumulation. T lymphocytes were stained with a panel of fluorescently labelled antibodies to simultaneously identify different T cell populations (MAIT, γδ, CD4 and CD8), and determine their activation status (CD69) and cytokine production (IFN-γ and TNF-α).

We first defined the heterogeneity of T cell responses to *Salmonella*, by performing an unsupervised clustering analysis on all CD3^+^ T cells expressing the activation marker CD69 following overnight incubation with the *Salmonella* pathovariants. Dimensionality reduction analysis by *t*-SNE revealed 22 populations of CD3^+^ CD69^+^ T cells, some of which differed in frequency according to the infecting *Salmonella* pathogen. Clusters were then annotated and assigned to MAIT (identified as CD3^+^ Vα7.2^+^ CD161^high^), γδ^+^, CD4^+^ or CD8^+^ T cell subsets based on the expression of distinct phenotypic markers.

We discovered that a group of clusters of MAIT cells (clusters 6, 15, 18, 21) was under-represented among all CD69^+^ cells upon infection with STM-D23580, compared with STM-LT2, ST-Ty2 and *E. coli* (Figure 1A). Next, we analysed expression of IFN-γ and TNF-α in CD69^+^ activated T cells. We determined that the under-represented clusters of CD69^+^ MAIT cells represented IFN-γ and TNF-α producing MAIT cells (Figure 1B). MAIT cells were next analysed using Uniform Manifold Approximation and Projection (UMAP), a neighbouring dimensionality reduction technique that preserves embedding and global distances better than t-SNE [39], and clearly defined the trajectory of the distinct subpopulations of *Salmonella*-activated MAIT cells (Figure 1C). STM-LT2 stimulated MAIT cells clustered close to MAIT cells stimulated with ST-Ty2 and *E. coli*, which were characterised by elevated expression of CD69 and the presence of single and double producers of TNF-α and IFN-γ. In contrast, STM-D23580 stimulated MAIT cells clustered closer to unstimulated cells, away from MAIT cells stimulated with ST-Ty2 and *E. coli* (Figure 1D, top left panel). STM-D23580 stimulated MAIT cells expressed low levels of CD69, with only a small TNF-α producing subpopulation and almost no IFN-γ producing cells (Figure 1D).

**Figure 1.**
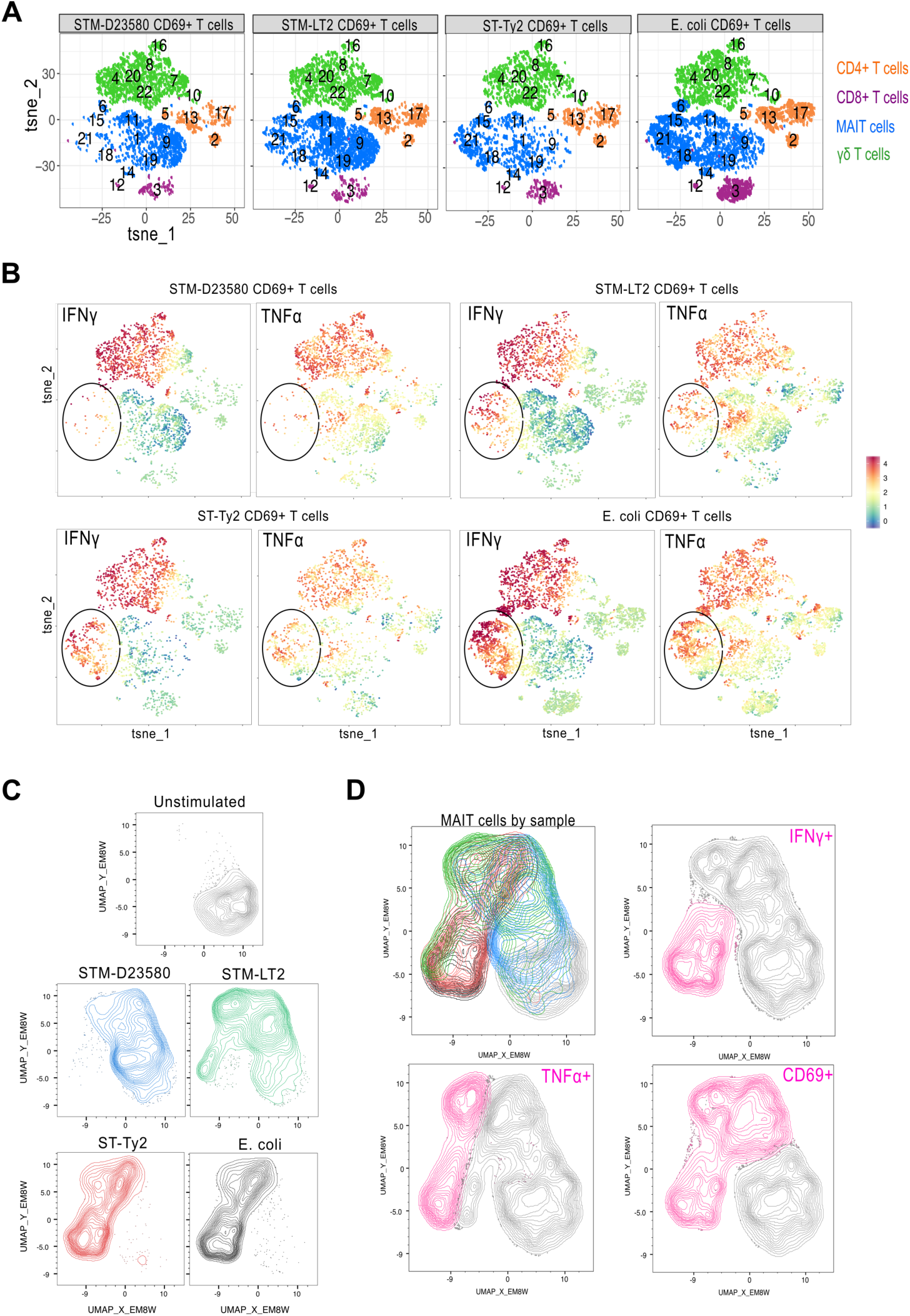
Identification of cellular responses to multiple *Salmonella enterica* subsp *enterica* serovars. PBMC were left unstimulated or were infected at MOI of 5 with STM-D23580, STM-LT2, ST-Ty2 or *E. coli*. Intracellular staining was performed to detect CD69 expression and cytokine production (TNF-α and IFN-γ), as correlates of T cell activation. **(A)** *t*-SNE plots on gated CD69^+^ CD3^+^ T cells infected with STM-D23580, STM-LT2, ST-Ty2 or *E. coli*. Four CD3^+^ T cell populations (CD4^+^, CD8^+^, γδ^+^ and MAIT) were annotated based on the expression of distinct phenotypic markers. CD4, CD8, TCRγδ, Vα7.2^+^, CD161, IFN-γ and TNF-α were the parameters included for *t*-SNE analysis. Plots correspond to one representative donor. **(B)** *t*-SNE plots as in (A) showing relative expression of TNF-α and IFN-γ on CD69^+^ CD3^+^ T cells. **(C)** UMAP analysis on concatenated CD3^+^ Vα7.2^+^ CD161^+^ MAIT cells from the same donor as in (A) and (B). Calculated UMAPs are shown for each experimental condition. CD69, IFN-γ and TNF-α were the parameters included for analysis. **(D)** Top left panel showing UMAP as an overlay of concatenated MAIT cell populations from (C): unstimulated in light grey, STM-D23580 in blue, STM-LT2 in green, ST-Ty2 in red and *E. coli* in dark grey. Top right and bottom panels: UMAPs showing expression of CD69, TNF-α and IFN-γ in pink. **(A-D)** Data from one donor representative of four biological replicates.

### *S.* Typhimurium sequence type 313 lineage 2 fails to elicit MAIT cell activation

To validate our unsupervised analysis, we infected PBMC with the different *Salmonella* strains at increasing multiplicity of infection (MOI), and then assessed MAIT cell activation by flow cytometry. Infection by STM-D23580 consistently induced limited MAIT cell responses across a range of MOIs and in every healthy donor tested. In comparison with STM-LT2, ST-Ty2 or *E. coli*, STM-D23580 stimulated MAIT cells significantly expressed less CD69 and produced less IFN-γ and TNF-α (Figure 2A-D). This effect was not dependent on a selective loss of MAIT cells, as STM-D23580 did not have a detrimental effect on MAIT cell viability (Figure S1A). γδ T cells, a subset of innate T lymphocytes also present in PBMC, were strongly activated by STM-D23580, indicating that the lack of cell activation is exclusive to MAIT cells (Fig 2A, 2C).

**Figure 2.**
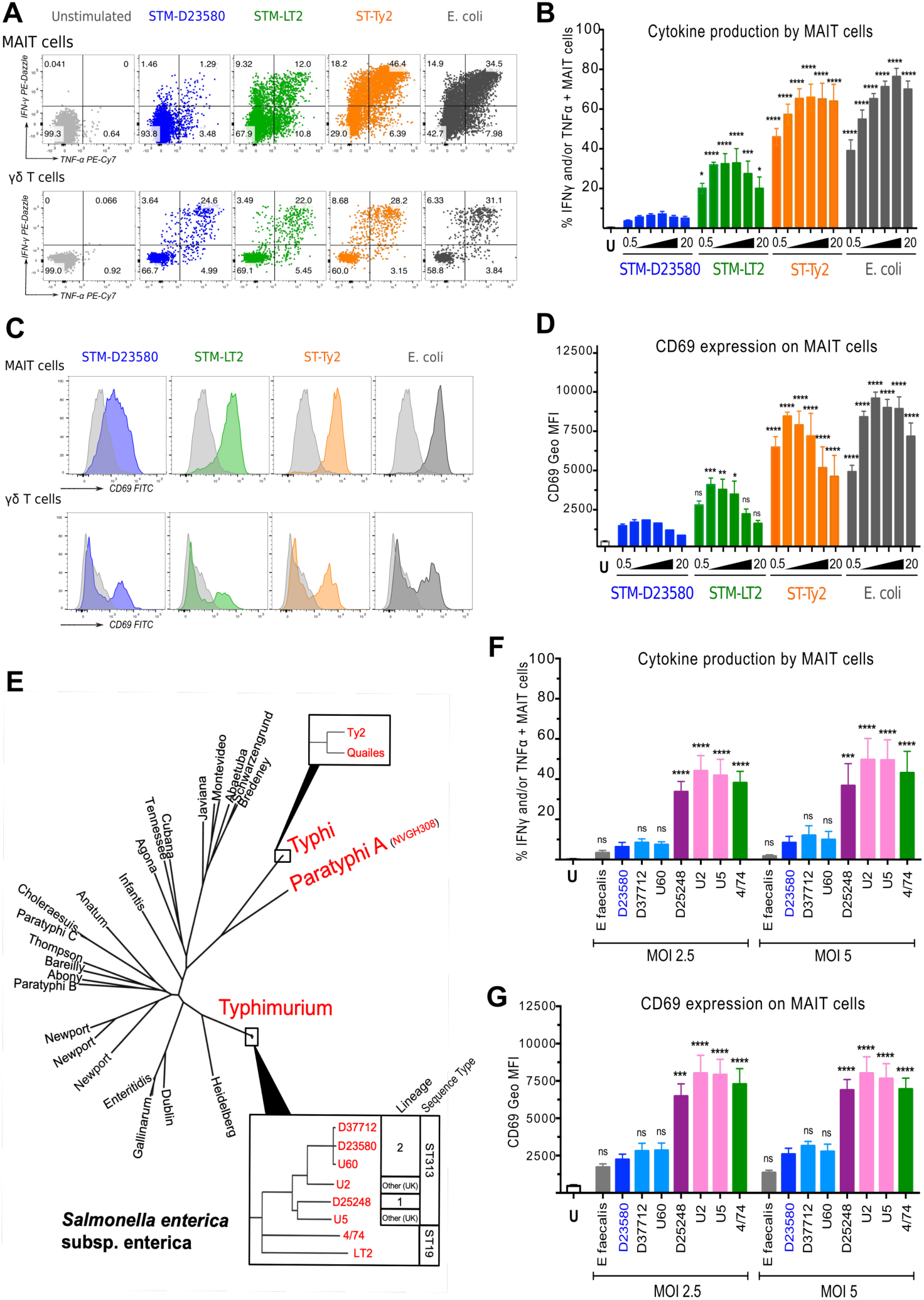
*S.* Typhimurium sequence type 313 lineage 2 fails to elicit MAIT cell activation. PBMC were left unstimulated (U) or were infected with a variety of *Salmonella* strains at the indicated MOI. *E coli* was included as positive control and *E. faecalis* as negative control. **(A)** Production of TNF-α and IFN-γ by MAIT and γδ^+^ T cells was detected by intracellular staining. Representative flow cytometry plots from one volunteer are shown. **(B)** Percentage of TNF-α and/or IFN-γ producing MAIT cells when stimulated at increasing MOI, from 0.5 to 20 bacteria per cell. Data represented as mean ± SEM, two-way ANOVA + Dunnet’s (vs. STM-D23580), n=4. **(C)** CD69 staining profile of stimulated MAIT and γδ^+^ T cells. Representative histograms from one volunteer are shown. **(D)** CD69 expression on MAIT cells when stimulated as in (B). Data represented as geometric mean ± SEM, two-way ANOVA + Dunnet’s (vs. STM-D23580), n=4. **(E)** Phylogenetic relationships between strains used in these experiments (red) within the context of *Salmonella enterica* phylogeny. **(F)** Percentage of TNF-α and/or IFN-γ producing MAIT cells, treated with bacterial strains at MOI of 2.5 and 5. Data represented as mean ± SEM, two-way ANOVA + Dunnet’s (vs. STM-D23580), n=4. **(G)** Levels of CD69 expression on MAIT cells treated as in (E). Data represented as geometric mean ± SEM, two-way ANOVA + Dunnet’s (vs. STM-D23580), n=4.

In culture, *Salmonella spp.* secrete vitamin B2 intermediates that can bind to MR1 on antigen presenting cells (APCs), to trigger MAIT cell activation [29]. To examine whether STM-D23580 secretes MAIT cell agonists, we collected supernatants from single-colony cultures to stimulate PBMC. Supernatants from STM-LT2 and *E. coli* induced a dose-dependent production of IFN-γ and TNF-α by MAIT cells, whereas STM-D23580 supernatants did not (Figure S1B).

To validate this finding, we examined MAIT cell responses to a broader selection of bacterial isolates, including two *S*. Typhi strains (ST-Ty2 and ST-Quailes) and one *S.* Paratyphi A strain; in addition two differently sourced stocks of STM-D23580 were tested, to ensure that genuine sequence type 313 isolates were being used. At two different MOIs, STM-D23580 elicited the lowest levels of MAIT cell activation of the group (Fig S1C-S1D). In contrast, γδ T cell responses were comparable across all *Salmonella* pathovars (S1E-S1F).

We next determined whether the lack of MAIT cell activation was caused by the entire *S.* Typhimurium sequence type 313 clade or was a unique characteristic of sequence type 313 lineage 2 which is currently causing most clinical disease in Africa [14]. STM-D23580 and additional isolates of sequence type 313 lineage 2, were compared with closely-related isolates that were members of sequence type 313 lineage 1 or the UK-sequence type 313 group that is associated with gastroenteritis (Figure 2E). To examine MR1-independent MAIT cell activation, we used *Enterococcus faecalis* as a negative control as it lacks the vitamin B2 biosynthetic pathway [29].

Remarkably, only the sequence type 313 strains belonging to lineage 2, such as D23580, D37712 and U60 failed to elicit MAIT cell activation (Figure 2F-2G). All other *Salmonella* sequence type 313 lineages triggered the same level of MAIT cell responses as *S*. Typhimurium 4/74 (STM-4/74) [40], a sequence type 19 strain that is closely related to STM-LT2 and is associated with non-invasive diarrhoeal infections.

### STM-D23580 does not affect MR1-dependent antigen presentation

To define the molecular mechanisms underlying the lack of MAIT stimulation by STM-D23580, we first investigated whether MAIT cell activation was MR1-dependent. Adding the anti-MR1 blocking antibody 26.5 [41] completely abrogated the MAIT cell activation induced by STM-LT2 and *E. coli*, as well as the minimal activation induced by STM-D23580 (Figure 3A-3B).

**Figure 3.**
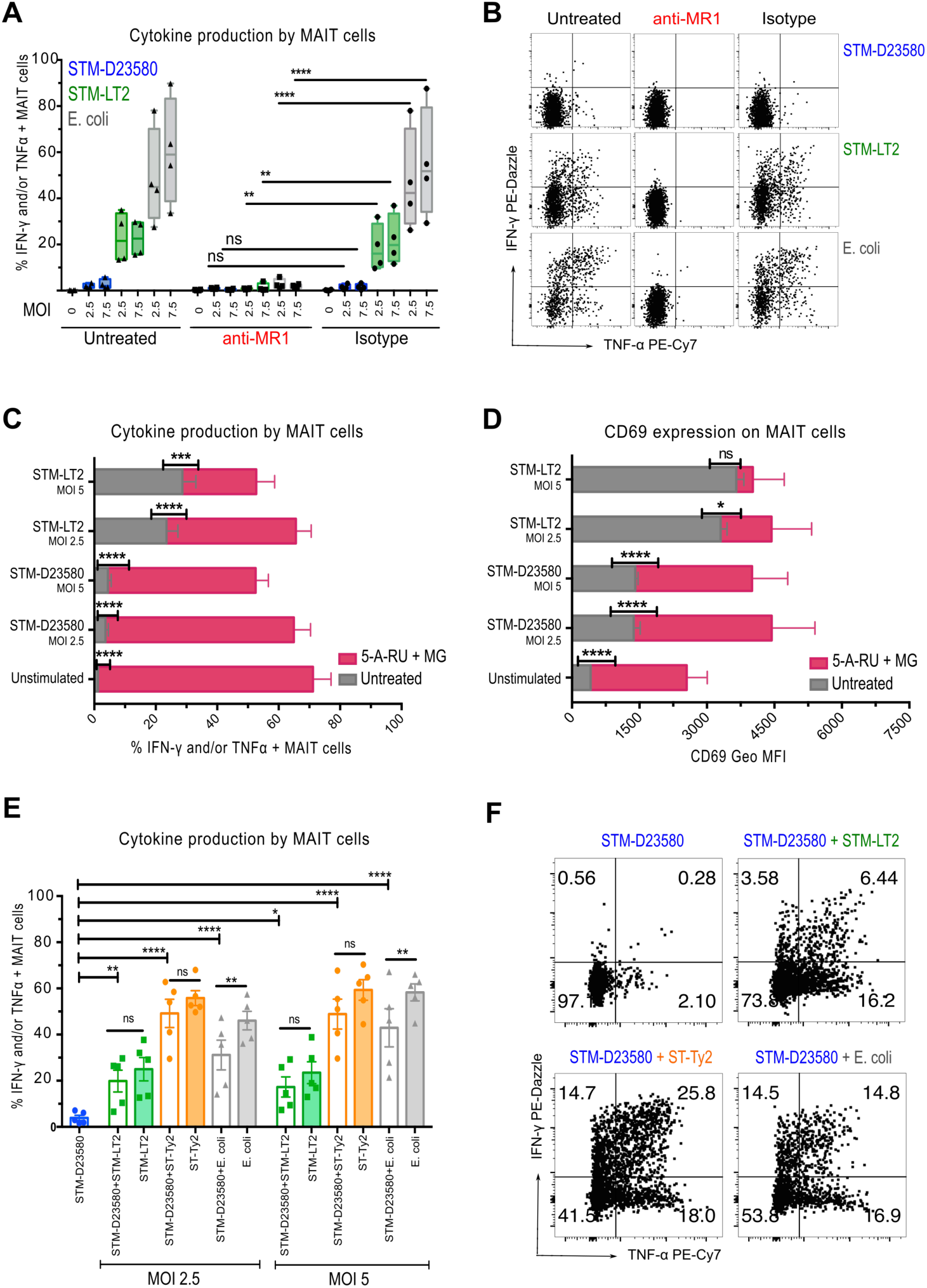
STM-D23580 does not affect MR1-dependent antigen presentation. **(A)** PBMC were infected (at MOI 2.5 and 7.5) with STM-D23580 (blue), STM-LT2 (green) or *E. coli* (grey), and incubated in the presence of anti-MR1 blocking antibody or the equivalent isotype control. Data are represented as percentage of TNF-α and/or IFN-γ producing MAIT cells, box-and-whisker plot, two-way ANOVA + Dunnet’s (vs. Isotype), n=4. **(B)** Representative example of cytokine production by stimulated MAIT cells treated as in (A). **(C-D)** PBMC were left unstimulated or were infected (at MOI 2.5 and 5) with either STM-D23580 or STM-LT2, in the presence (pink bars) or absence (grey bars) of the MR1 ligands 5-A-RU and MG. Percentage of cytokine producing MAIT cells and their CD69 expression are shown. Data represented as mean and geometric mean ± SEM, two-way ANOVA + Bonferroni’s, n=4. **(E)** PBMC were infected with D23580 at MOI of 2.5, alone or in combination with STM-LT2 (green), ST-Ty2 (orange) or *E. coli* (grey), at two different MOI (2.5 and 5). Data represented as percentage of TNF-α and/or IFN-γ producing MAIT cells, mean ± SEM, one-way ANOVA + Sidak’s, n=5. **(F)** Representative example of cytokine production by stimulated MAIT cells treated as in (E).

We next examined whether STM-D23580 either failed to produce stimulatory MR1 ligands or actively inhibited MAIT cell activation. MAIT cell activation was restored following the addition of the canonical MAIT cell ligand 5-amino-6-D-ribitylaminouracil (5-A-RU) and methylglyoxal (MG) [33] to infected PBMC (Figure 3C-3D), demonstrating that a dominant antagonistic MR1 ligand was not released by STM-D23580. Consistent with these results, a combination of supernatants from overnight cultures of both STM-D23580 and STM-LT2 (added simultaneously or one hour apart) fully restored MAIT cell activation (Figure S2A-2SB). Likewise, co-infection of PBMC with STM-D23580 plus either STM-LT2, ST-Ty2 or *E. coli* also restored MAIT cell activation to the levels observed with single infections (Figure 3E-3F). Taken together, our data show that STM-D23580 neither interferes nor blocks MAIT cell activation in the presence of stimulatory MR1 ligands.

To exclude the possibility that the lack of MAIT cell activation arose from an insufficient infection of APCs, we exposed Monocyte-derived Dendritic Cells (MoDCs) to fluorescently labelled STM-D23580 or STM-LT2. Live *Salmonella*-containing MoDCs were FACS-sorted and co-cultured with enriched autologous CD3^+^ T lymphocytes, as described previously [42]. In contrast to STM-LT2 infected MoDCs, STM-D23580 infected MoDCs did not stimulate effector MAIT cells (Figure S2C). Using an MR1-overexpressing antigen presenting cell line, we also excluded downregulation of surface MR1 expression in the presence of STM-D23580 supernatants as a possible cause of the lack of MAIT cell activation (Figure S2D).

Cytokines, such as IL-12, IL-18 and type I IFN, released by APCs upon bacterial or viral infection can also activate MAIT cells in a MR1-independent manner [31]. We confirmed that when MoDCs were co-cultured with purified MAIT cells, equal amounts of bioactive IL-12p70 were secreted upon infection with STM-D23580, STM-LT2 and *E. coli* (Figure S2E).

Taken together, these observations refuted the hypothesis that impaired MR1-dependent antigen-presentation followed STM-D23580 infection.

### STM-D23580 evades MAIT cell recognition by overexpression of RibB

Our results suggest that STM-D23580 and related isolates from the sequence type 313 lineage 2 might not produce the MR1 binding ligands that are generated by other *S.* Typhimurium or *S.* Typhi pathovariants. The major source of natural antigens driving MAIT cell activation derives from by-products of microbial riboflavin synthesis [33]. We depict the *Salmonella* riboflavin biosynthetic pathway in Figure 4A. A comparison of the coding sequences (CDS) of the enzymes involved in the riboflavin biosynthesis pathway found only one nucleotide change (SNP) between the sequence type 19 STM-4/74 and the sequence type 313 STM-D23580 strains. This synonymous coding variant was located in the *ribD* gene (Glu316Glu) [43], suggesting that there were no biochemical differences between the riboflavin biosynthesis pathways of the sequence type 313 and sequence type 19 isolates.

**Figure 4.**
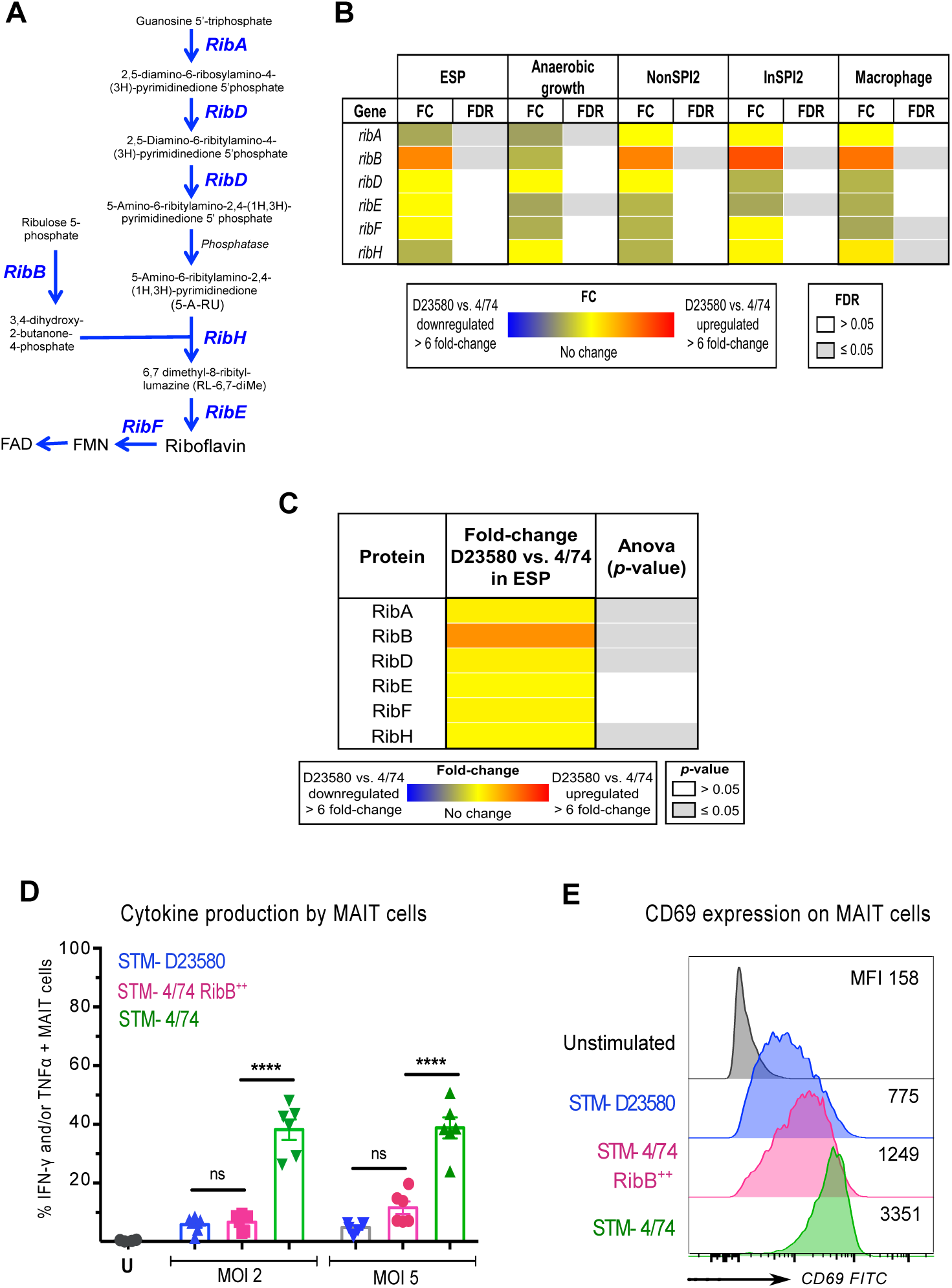
STM-D23580 evades MAIT cell recognition by overexpression of *ribB*. (**A**) Schematic representation of *Salmonella*’s riboflavin pathway, adapted from Soudais *et al*. [52]. **(B)** Heat map of the *rib* genes from RNA-seq data sets obtained from STM-D23580 and STM-4/74 in five infection-relevant conditions: ESP (early stationary phase), anaerobic growth, NonSPI2 (SPI2-non-inducing PCN), InSPI2 (SPI2-inducing PCN), and macrophage (intra-RAW264.7 murine macrophage environment). Values indicate fold-change (FC) and false discovery rate (FDR) calculated performing a Voom/Limma analysis (using Degust) for the RNA-seq data comparison of STM-D23580 versus STM-4/74. Data represent three biological replicates. PCN = phosphate carbon nitrogen minimal medium. **(C)** Heat map of the differential expression analysis at the protein level of the Rib proteins in the ESP condition. Results from LC-MS/MS were analysed using the Progenesis QI software (Nonlinear Dynamics) for label-free quantification analysis. Each sample represents six biological replicates. Data represented in panels **B** and **C** were extracted from Canals *et.al.* [43]. **(D)** PBMC were infected at two different MOI (2 and 5) with STM-D23580 (blue), STM-4/74 (green) or STM-4/74 RibB^++^ (pink). Data represented as percentage of TNF-α and/or IFN-γ producing MAIT cells, mean ± SEM, two-way ANOVA + Tukey’s, n=6. **(E)** CD69 staining profile of stimulated MAIT cells treated as in (D). Representative histograms from one volunteer are shown, MFI=Median Fluorescence Intensity.

To determine whether the enzymes of the riboflavin pathway of the sequence type 19 STM-4/74 and sequence type 313 STM-D23580 strains were expressed at different levels, we investigated the transcriptomic and proteomic data from our recent comparative analysis [43]. Strains STM-4/74 and STM-D23580 are closely-related, sharing 92% of coding genes [43]. Differential gene expression analysis of the *rib* genes at the transcriptomic level identified significant up-regulation of *ribB* (≥2 fold-change, FDR ≤0.001) in STM-D23580 in four out of the five experimental conditions (Figure 4B). We then examined data from a quantitative proteomic approach which showed that RibB protein levels were up-regulated in STM-D23580 compared to STM-4/74, during growth in rich medium at early stationary phase (ESP) (Figure 4C).

We searched for a molecular explanation for the high levels of *ribB* expression in STM-D23580 compared to STM-4/74. In STM-D23580, *ribB* and its 5’ untranslated region (5’UTR) are transcribed as a single transcript that is initiated from a single gene promoter which we identified previously [43]. By analogy with the genetic mechanism identified for over-expression of the PgtE virulence factor in STM-D23580 [44] we searched for nucleotide polymorphisms that distinguished the *ribB* regions of the two strains. There were no differences between the promoter sequences of the *ribB* genes or the 5’ UTR of the strains STM-D23580 and STM-4/74.

To determine whether the MAIT cell activation phenotype was linked to the overexpression of the RibB enzyme (4-dihydroxy-2-butanone 4-phosphate synthase), we created a derivative of STM-4/74 that expressed high levels of RibB. Since deletions in the riboflavin biosynthetic pathway genes are lethal without high dose riboflavin supplementation [33][45] and *ribB* is essential for *Salmonella in vivo* virulence [46], we used a gene cloning approach to overexpress the *ribB* gene of STM-D23580 in STM-4/74 from a recombinant plasmid (STM-4/74 RibB^++^). The expression of high levels of the RibB enzyme by STM-4/74 ablated the substantial MAIT cell activation induced by wild type STM-4/74. Infection with STM-4/74 RibB^++^ induced low levels of cytokine production and CD69 expression by MAIT cells, recapitulating the phenotype of STM-D23580 (Figure 4D-4E). Importantly, γδ T cells responded equally to both STM-4/74 wild type and STM-4/74 RibB^++^ (Figure S3).

To investigate whether overexpression of RibB altered the balance of downstream products from the riboflavin pathway, we measured the amount of riboflavin, flavin mononucleotide (FMN) and flavin adenine dinucleotide (FAD) by HPLC. Following growth to early stationary phase, intracellular samples and culture supernatants from STM-4/74 and STM-4/74 RibB^++^ contained larger amounts of riboflavin than the sequence type 313 strains STM-D23580 and STM-D37712 (Figure 5A-5B). The STM-4/74 RibB^++^ supernatant with the highest level of riboflavin also contained the largest amount of FMN, compared with STM-4/74, STM-D23580 and STM-D37712 (Figure 5C).

**Figure 5.**
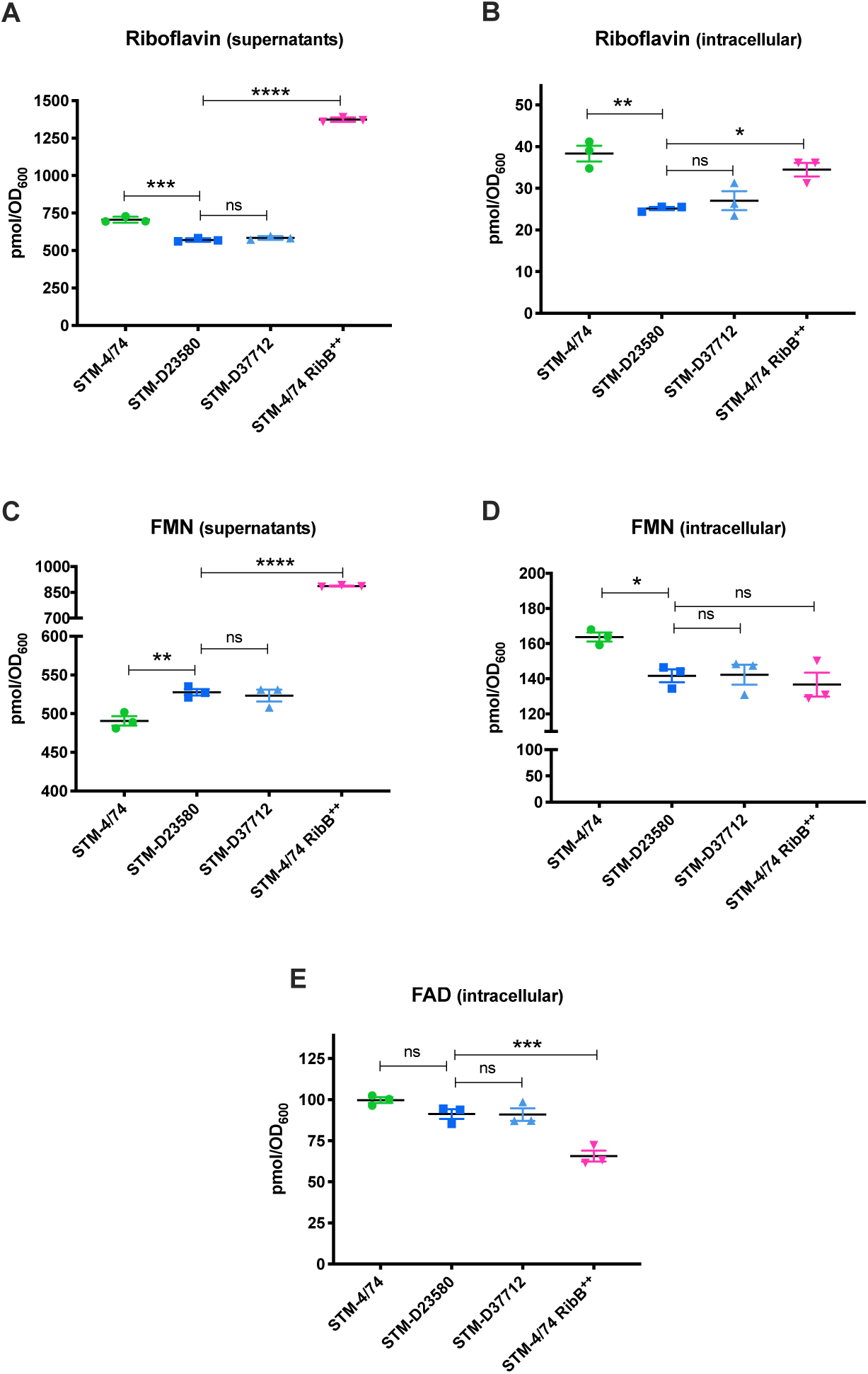
STM-4/74 RibB^++^ has the lowest intracellular levels of FMN, a negative regulator of *ribB* gene expression. Supernatants from early stationary phase and bacterial pellets were harvested and analysed by HLPC using riboflavin, FMN and FAD standards. Data is reported in pmol and has been normalised to the absorbance (OD_600_) from each culture. **(A)** Riboflavin levels in supernatants. **(B)** Intracellular riboflavin levels. **(C)** FMN levels in supernatants. **(D)** Intracellular FMN levels. **(D)** Intracellular FAD levels. Measurements were obtained from 3 biological replicates, mean ± SEM, one-way ANOVA + Dunnet’s (vs. STM-D23580).

Overall, the data show that the RibB over-producing strain (4/74 RibB^++^) produced the highest level of extracellular riboflavin and FMN, and had the lowest level of intracellular FMN. In contrast, the STM-4/74 wild-type strain had the largest intracellular amount of FMN, compared with 4/74 RibB^++^ and both of the sequence type 313 strains (Figure 5D). While intracellular FAD levels were similar between STM-4/74, STM-D23580 and STM-D37712, STM-4/74 RibB^++^ contained the lowest level of intracellular FAD (Figure 5E). It was not possible to measure the levels of MAIT activator ligands such as 5-OP-RU (5- (2-oxopropylideneamino)-6-D-ribitylaminouracil) or 5-OE-RU (5- (2-oxoethylideneamino)-6-D-ribitylaminouracil), products of the riboflavin pathway, due to the unstable nature of these metabolites [33]. We conclude that the lowest levels of intracellular FMN was found in the STM-D23580 and STM-D37712, STM-4/74 RibB^++^ strains that overexpress the RibB enyzyme.

### Characterising MAIT cell responses to *Salmonella* in an immunocompromised patient cohort

To validate our observations in a clinically relevant population, we performed a subset of our assays on blood samples obtained from HIV-infected adults living in Malawi, an endemic area for iNTS infections with sequence type 313 strains. Comparing a cohort of healthy samples from the UK and Malawi, and consistent with previous reports (reviewed in [47]), the overall percentage of MAIT cells was reduced among HIV-infected individuals, particularly those not receiving antiretroviral therapy (ART) (Figure 6A). In line with our previous data from healthy volunteers, MAIT cells from HIV-infected individuals failed to produce IFN-γ and TNF-α following *ex vivo* stimulation with STM-D23580, regardless of their ART status (Figure 6B). However, MAIT cells from HIV^+^ adults responded strongly to *Salmonella spp*. including *S.* Typhi and *S.* Typhimurium STM-4/74. Importantly, this result suggests that evasion of MAIT cell recognition by sequence type 313 strains could be a critical factor during the course of natural iNTS disease in HIV-infected individuals.

**Figure 6.**
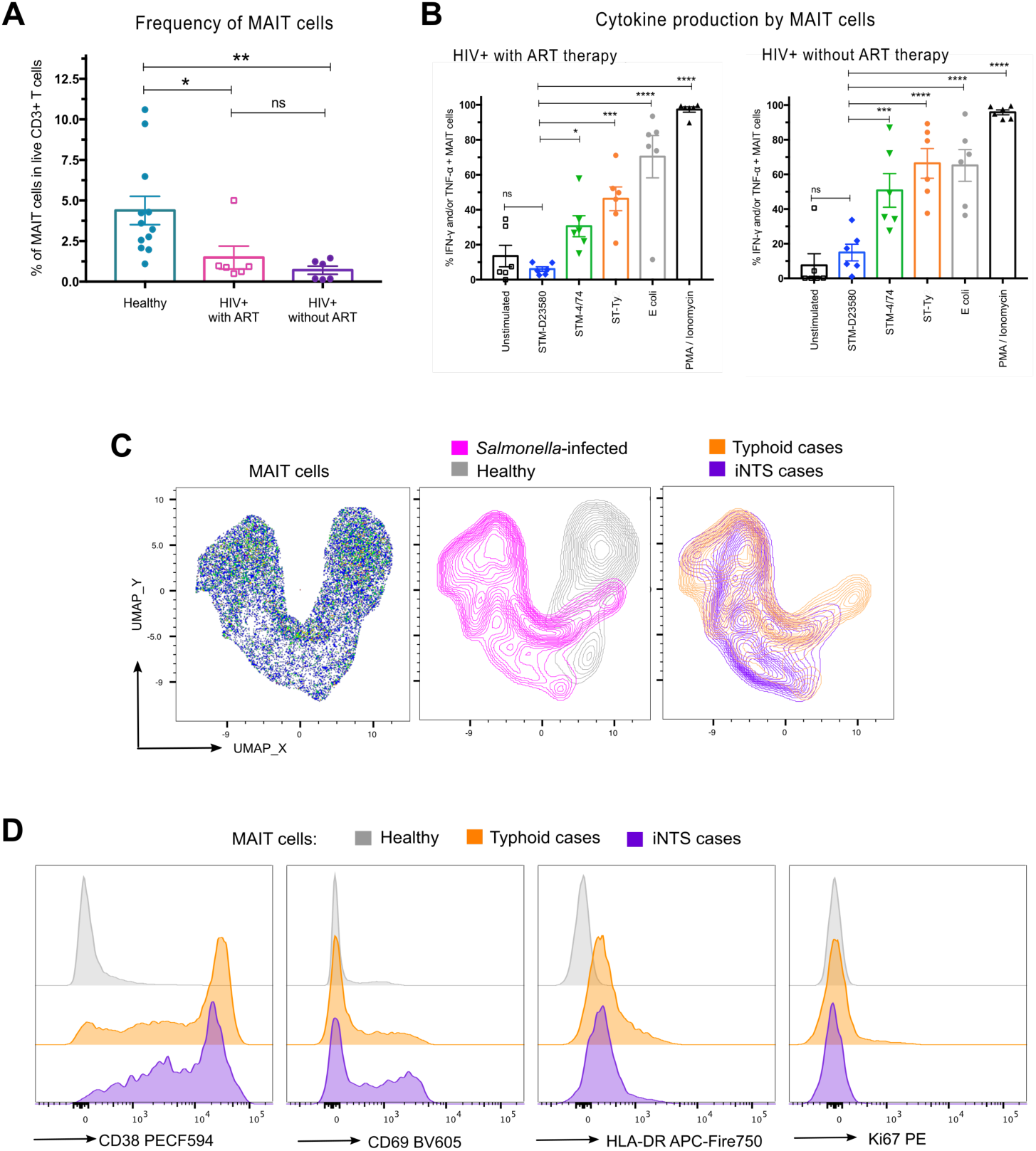
Characterising MAIT cell responses to *Salmonella* in an immunocompromised patient cohort. **(A)** Percentage of MAIT cells (CD3^+^ Vα7.2^+^ CD161^+^) in PBMC isolated from healthy (n=12) and HIV-infected patients with (n=6) or without ART (n=6). Data represented as percentage of live CD3^+^ T cells, mean ± SEM, Kruskal-Wallis + Dunn’s. **(B)** PBMC isolated from HIV^+^ patients with or without ART were infected at MOI of 7 with either STM-D23580, STM-4/74, STy-H58 or *E. coli*. PMA/ionomycin was used as positive control. Intracellular staining was performed to detect cytokine production (IFN-γ and TNF-α) as correlate of MAIT cell activation. Data represented as percentage of TNF-α and/or IFN-γ producing MAIT cells, mean ± SEM, one-way ANOVA + Dunnet’s (vs. STM-D23580), n=6 for each group of patients. **(C)** UMAP analysis was performed on concatenated manually gated MAIT cells from cases and healthy controls. The distribution of iNTS and typhoid cases is shown as overlay contour plots. **(D)** Histograms showing expression of activation markers (CD38, CD69, HLA-DR, Ki67) on concatenated MAIT cells from (D).

To capture MAIT cells activity *in vivo* during natural infection we examined fixed/lysed whole blood aliquots from *Salmonella* infected patients for markers of T cell activation. Blood samples were obtained from five typhoidal and three iNTS cases at Queen Elizabeth Central Hospital (Blantyre, Malawi), shortly after diagnosis by blood culture (range 2-9 days). The proportions of the major T cell subsets differed significantly between typhoidal and nontyphoidal cases (Figure S4A). The proportion of CD4^+^ T cells in particular was significantly reduced in nontyphoidal cases compared with controls, most likely as a result of the underlying HIV infection. We observed a compensatory enlargement of the percentage of CD8^+^ T cells which was not accompanied with changes in the proportion of MAIT cells (Figure S4A).

We concatenated CD3^+^ T cells and MAIT cells from cases and controls, and analysed by dimensionality reduction using UMAP, including all phenotypic and activation parameters (Figure S4B and Figure 6C). UMAP of MAIT cells identified and clustered distinct Salmonellosis cases from healthy individuals, but did not differentiate between iNTS and typhoid cases (Figure 6C). This likely reflects heterogeneity in the small group of subjects, particularly variance in time courses and clinical pathogenesis of iNTS disease compared to typhoid, and the confounding effect of antibiotic treatment in clinical disease. Histograms on concatenated MAIT cells by group, confirmed a significant induction of CD38, CD69 and HLA-DR upregulation in *Salmonella*-infected individuals when compared with healthy controls (Figure 6D and Figure S5). Percentages of proliferating MAIT cells, identified based on Ki67 intracellular expression, varied significantly among individuals in this small sample (Figure S5).

## DISCUSSION

Here we investigated the ability of MAIT cells to recognise and respond to diverse invasive *Salmonella enterica* serovars. We found that MAIT cells isolated from the blood of healthy and HIV-infected individuals were not activated by exposure to invasive disease-associated *S.* Typhimurium sequence type 313 lineage 2 strains. Our data demonstrated how *S.* Typhimurium sequence type 313 lineage 2 evades MAIT cell recognition by overexpressing *ribB*, a bacterial gene encoding the RibB enzyme involved in the riboflavin pathway. Our results lead us to propose that this RibB-mediated mechanism provides an evolutionary advantage that allows invasive *S.* Typhimurium sequence type 313 lineage 2 bacteria to escape cell immune responses by overexpressing a single riboflavin bacterial gene.

MR1-restricted MAIT cells are highly abundant in the gut mucosa [48], where they reach their final maturation upon recognition of vitamin B2 metabolites derived from gut commensals presented by MR1 expressing mucosal B cells [48]. MAIT cells show antimicrobial activity *in vivo* and *in vitro* [49], [50] through MR1 dependent and independent interactions. Microorganisms must express the riboflavin biosynthesis pathway to be able to activate MAIT cells [29], [51]. Mutations in key enzymes of the riboflavin biosynthetic pathway in both Gram positive and negative bacteria can abrogate MAIT cell activation [33], [52]. Different bacteria that possess the riboflavin biosynthetic pathway induce varying levels of MAIT stimulation [36], [37], possibly through the influences of the microenvironment on bacterial metabolism and antigen availability, or the known short half-life of the potent MAIT cell antigens, 5-OP-RU and 5-OE-RU [53]. The ability of MAIT cells to recognise and respond to several isolates of the same pathogen may also vary to reflect bacterial metabolic differences. For example, riboflavin metabolism variation among clinical isolates of *Streptococcus pneumoniae* produces different measurable levels of riboflavin and FMN that correlate with differential activation of MAIT cells [38].

*Salmonella* spp. possess an active riboflavin biosynthetic pathway, which generates MAIT cell agonists [29], [33]. MAIT cells recognise and kill *S.* Typhimurium-infected targets *in vitro* [32], and activated MAIT cells accumulate in murine lungs following intranasal infection with *S.* Typhimurium [26]. However, bacterial lung clearance was independent of MAIT cells in this infection model, possibly due to the non-physiological route of infection. Human studies demonstrated sustained MAIT cell activation and proliferation at the peak of infection with *S.* Typhi and *S.* Paratyphi [34], [35]. Among immunocompetent humans, the clinical outcomes of infection by *Salmonella enterica* spp. depend on the infecting serovar. Human-restricted typhoidal serovars, such as *S*. Typhi induce the most severe form of systemic disease, typhoid fever; while the broad-host *S.* Typhimurium sequence type 19 causes self-limiting gastroenteritis. The recently documented multidrug resistant *S.* Typhimurium sequence type 313 clade causes the majority of iNTS cases among immunocompromised adults and malnourished young children living in sub-Saharan Africa.

Several bacterial factors have been reported to enhance invasiveness of *S*. Typhimurium sequence type 313, suggesting a multifactorial adaptation of this African lineage to a systemic lifestyle. These include interference with the complement cascade [44], interference with DC function [42], reduced inflammasome activation [54], dissemination through CD11b^+^ migratory DC [55] among others.

Here we found that *S.* Typhi, *S*. Paratyphi A and most *S.* Typhimurium pathovars potently elicited *ex vivo* MR1-dependent MAIT cell activation, but all tested isolates from the invasive *S.* Typhimurium sequence type 313 lineage 2 barely induced cytokine secretion or CD69 upregulation. This suboptimal activation was restricted to MAIT cells, as γδ cell activation was comparable across all isolates tested. Our data excludes differences in infection efficiency, MR1 expression, co-stimulatory cytokines (IL-12), the genetic sequence of the riboflavin encoding enzymes of *S.* Typhimurium and the presence of dominant inhibitory ligands as potential mechanisms for these results. Following comparative proteomic and transcriptomic analyses, we discovered that invasive *S.* Typhimurium sequence type 313 lineage 2 pathovars escape MAIT cell recognition by overexpressing *ribB*, a bacterial gene encoding the riboflavin biosynthetic enzyme RibB. By overexpressing this single riboflavin gene in a sequence type 19 *S*. Typhimurium strain, we revealed that up-regulation of this single riboflavin gene was sufficient to abrogate MAIT cell responses.

Transcriptional control of the bacterial *ribB* gene is controlled by a conserved FMN riboswitch, which is located in the 5′ untranslated region (5’ UTR) of *ribB* [56], and is negatively regulated by FMN and other flavins at the transcriptional and translational levels in *E. coli* [57]. The SroG small RNA (sRNA) is derived from the *ribB* 5’ leader sequence, although the function of this sRNA remains unknown [58]. In addition to the riboswitch-mediated regulation, RibB expression is induced by growth in a low pH environment [59]. In STM-D23580, both *sroG* and *ribB* are transcribed as a single transcript that is initiated from a single gene promoter which we identified previously [43].

While the precise molecular mechanism responsible for *ribB* overexpression remains to be established, a single noncoding nucleotide polymorphism in the promoter of the *pgtE* gene of the sequence type 313 lineage 2 strain STM-D23580 is known to be responsible for high expression of the outer membrane PgtE virulence factor, which promotes bacterial survival and dissemination during mammalian infection [44]. The lack of nucleotide differences between the promoter sequences of the *ribB* genes or the riboswitch of the two strains suggests that the high level of expression of *ribB* in STM-D23580 is caused by a novel and uncharacterised regulatory mechanism.

It has been proposed that genomic changes in sequence type 313 isolates that confer altered metabolism and increased anaerobic metabolic capacity are linked to adaptation of the extra-intestinal niche [14]. Riboflavin and its derivatives are important cofactors for flavoproteins involved in cellular redox metabolism and several biochemical pathways, proposed to be essential for the metabolic adaptation of the sequence type 313 clade. Riboflavin promotes intracellular microbial survival and virulence during *in vivo* infection with *Histoplasma capsulatum* and *Brucella abortus* [60], [61]. In addition, accumulation of riboflavin is a candidate virulence factor in *Pseudogymnoascus destructans* skin infection [62].

The metabolomic measurements of the end-products of the riboflavin pathway showed a correlation between lower levels of intracellular FMN and increased expression of RibB. Because FMN is a negative regulator of *ribB* gene expression [57], we speculate that the lower levels of intracellular FMN observed in the ST313 strains D23580 and D37712 are linked to the high levels of expression of RibB in these African *S.* Typhimurium strains.

We acknowledge that our *in vivo* validation was limited by the small sample size of iNTS cases that could be obtained, as well as by the inherently heterogeneous characteristics of disease presentation for typhoid fever compared to iNTS disease. For example, our cohort of *Salmonella*-infected patients recruited at one site in Malawi showed a wide variability in clinical presentation, including the time between symptoms and receiving medical attention and the time between a positive blood culture and sample collection. These factors plus co-morbidities, such as HIV and tuberculosis, decreased the power of our iNTS disease cohort. While some mouse models of infection with *Salmonella* sequence type 313 strains have been published [14], [63], these models present limited utility as mice have very low frequencies and absolute numbers of MAIT cells as compared to humans [28]. Our findings may be of major relevance during the initial phase of infection, in the gut, where is expected that resident immune cells such as MAIT cells should prevent systemic infection by encountering and responding rapidly to bacterial signals.

Overall, our findings suggest that MAIT cells play a crucial role in defence against invasive *Salmonella* disease in humans and that evasion from MAIT cell recognition is a critical mechanism for the invasiveness of *S*. Typhimurium sequence type 313 lineage 2 isolates. Our results propose, for the first time, how differences in MAIT cells activation may associate to distinct diseases caused by closely related microorganisms. The increased susceptibility of immunocompromised patients to the *S*. Typhimurium sequence type 313 lineage 2 strains suggest that MAIT cells might play a particularly relevant role in the context of waning CD4^+^ T cell mediated protective adaptive immunity. For example, following HIV co-infection and/or malnutrition, among individuals suffering from recurrent gut infections secondary to intestinal barrier dysfunction [64], [65], microbiota dysbiosis [66], [67] and multiple innate and adaptive immune defects [68]. We propose that the ability of MAIT cells to target gastrointestinal pathogens represents a key immunological evolutionary bottleneck that has been effectively countered by *Salmonella*, resulting in the current epidemic of invasive disease in Africa.

## METHODS

### Bacterial strains and preparation of stocks

This study included representative strains of *Salmonella enterica serovar* Typhimurium, from both the sequence type 19 and the sequence type 313. *S.* Typhi and *S.* Paratyphi A serovars were utilised as comparative Typhoidal invasive strains, while *Escherichia coli* (DH5α) was used as unrelated bacterial control. Supplementary Table 1 lists bacterial strains used in this study.

Overnight bacterial cultures from a single colony origin were used to inoculate LB Lennox broth (Sigma) supplemented with sucrose (Sigma) at a final concentration of 10%. Inoculated cultures were incubated at 37°C under constant shaking for approximately 3 hours, until reaching mid logarithmic phase. Bacterial aliquots were prepared and immediately frozen at −80°C for long-term storage. Bacterial viability of frozen aliquots was monitored periodically in order to maintain experimental reproducibility. The number of viable colony forming units (CFU) was determined with the Miles and Misra method, by plating 10-fold dilutions of the bacterial suspension onto LB Lennox agar (Sigma). On the day of the experiment, a single aliquot was thawed, washed twice with PBS and re-suspended in RPMI 1640 media to obtain the desired multiplicity of infection (MOI).

In the case of bacterial supernatants, these were taken from late exponential phase cultures, grown from a single colony following 18 hours incubation under constant shaking. Supernatants were filter sterilised before using.

### Construction of 4/74 pP_L_-*ribB*

To construct pP_L_-*ribB*, the *ribB* gene was amplified from genomic DNA of *S.* Typhimurium 4/74 using primers *ribB*_FW and *ribB*_RV. The PCR product was used for a linear amplification reaction with plasmid pJV300 (pP_L_) using Phusion DNA polymerase (New England Biolabs), and the resulting product was digested with DpnI. The plasmid was transformed into *E. coli* TOP10 and selected on LB plates supplemented with 100 μg/mL ampicillin. Plasmid presence was confirmed by PCR and DNA sequencing using oligonucleotides pPL_Seq_FW and pPL_Seq_RV. The pP_L_-*ribB* plasmid was subsequently purified and transformed into *S.* Typhimurium 4/74. Supplementary Table 2 lists plasmids and oligonucleotides used in this study.

### Isolation of human cells from peripheral blood from healthy volunteers in the UK

Leukocyte Reduction System cones were obtained from the UK National Blood Centre with informed consent following local ethical guidelines (NHSTB account T293). Blood was diluted in PBS and separated by gradient centrifugation using Lymphoprep™ (AxisShield). Peripheral Blood Mononuclear Cells (PBMC) were collected from the interface, washed with PBS, resuspended in complete medium and counted. Complete medium used throughout was RPMI 1640 (Sigma), supplemented with 10% heat-inactivated FCS (Sigma), 2 mM L-glutamine, 1% nonessential amino acids and 1% sodium pyruvate (all from Gibco).

### Isolation of PBMC from *Salmonella*-infected patients and HIV-infected patients

Blood samples from *Salmonella* infected adults were obtained as part of a case-control study carried out at Queen Elizabeth Central Hospital (Blantyre, Malawi) and conducted following local ethical guidelines (P09/17/2284). Potential study participants, with a history of fever, were identified and recruited only after *Salmonella* infection was confirmed by a blood culture taken as part of their routine clinical care. After obtaining appropriate consent and medical authorisation, a peripheral blood sample was collected between days 2 and 9 after positive blood culture. Aliquots of whole blood were treated with Fix/lyse buffers (SmartTube) and stored frozen until use. For analysis, cases were divided in invasive nontyphoidal (iNTS) or typhoidal Salmonellosis based on microbiology testing. As part of the same study, healthy adults from the local community were enrolled as controls. A peripheral blood sample was obtained from these individuals and processed in the same way as the cases. Since iNTS cases occur in immunosuppressed individuals, HIV+ patients were enrolled as extra controls. Adults presenting for HIV testing at the voluntary testing clinic, the HIV outpatient clinic, and the medical inpatient wards at the Queen Elizabeth Central Hospital were recruited. Based on the use of antiretroviral therapy, these patients were classified as ART naïve (without) or ART treated (with). Upon appropriate consent and medical authorisation, a blood sample was collected and PBMC were isolated for *ex vivo* infection assays, as described below.

### *Ex vivo* infection assays with PBMC

PBMC were seeded in 96 well plates round bottom (5 to 8 ×10^5^ cells per well) and infected with the different *Salmonella* strains from frozen mid-log phase stocks, at the indicated MOI. Upon 80 minutes incubation at 37°C, 100 μg/mL gentamicin (Lonza) was added to kill extracellular bacteria. At 180 minutes post-infection, 5ng/mL brefeldin A (BioLegend) solution was added to every well in order to achieve accumulation of intracellular cytokines. Samples were incubated overnight at 37°C for no more than 15 hours.

For MR1 blocking experiments, infection was performed in the presence of 30 μg/mL MR1 blocking antibody 26.5 [41] or mouse IgG2a isotype control (ATCC). The MAIT agonists 5-Amino-6-D-ribitylaminouracil (5-A-RU) was synthesised as described [69] and 1 μg/mL was combined to 50 μM methylglyoxal (MG, Sigma).

### Assessment of cytokine production and MAIT cell activation by flow cytometry

Following incubation in the presence of brefeldin A, cells were harvested, washed and stained with a viability dye (live/dead Zombie Aqua, BioLegend) for 20 minutes. Fixation was performed for 30 minutes at 4°C using the Foxp3 Fixation/Permeabilization buffers (eBioscience). Fixed cells were permeabilised and stained with an antibody cocktail for 40 minutes at room temperature, washed and stored protected from light at 4°C in PBS with 0.5% bovine serum albumin (FACS buffer) until acquisition. The following antibodies were used for extracellular and intracellular staining as 2 different panels: anti-CD3 Alexa700 (clone UCHT1; BioLegend), anti-CD3 PerCp Cy5.5 (clone UCHT1; BioLegend), anti-CD4 APCef780 (clone RPA-T4; eBioscience), anti-CD4 Alexa700 (clone RPA-T4; BioLegend), anti-CD8 BV785 (clone RPA-T8; BioLegend), anti-TCR γ/δ APC (clone B1; BioLegend), anti-CD161 BV605 (clone HP-3G10; BioLegend), anti-CD161 BV421 (clone HP-3G10; BioLegend), anti-Vα7.2 PE (clone 3C10; BioLegend), anti-Vα7.2 PE-Cy7 (clone 3C10; BioLegend), anti-CD69 FITC (clone FN50; BioLegend), anti TNF-α PECy7 (clone Mab11; BioLegend), anti TNF-α APC (clone Mab11; BioLegend), anti-IFN-γ FITC (clone 4S.B3; BioLegend) and anti-IFN-γ PE Dazzle (clone 4S.B3; BioLegend). Samples from the UK were acquired on a FortessaX20 (BD), whilst samples from the case-control study in Malawi were acquired on a LSR Fortessa cytometer (BD). All data was analysed with the same gating strategy on FlowJo (v.10.4.1).

### Assessment of immune profile in fixed/lysed whole blood samples by flow cytometry

One aliquot of fixed/lysed whole blood was defrosted and washed three times in FACS buffer. Cells were stained for surface and intracellular markers as described above and acquired on a LSR Fortessa cytometer. The following antibodies were included in one cocktail containing brilliant stain buffer (BD Biosciences): anti-CD3 PerCp Cy5.5 (clone UCHT1; BioLegend), anti-CD4 BV510 (clone RPA-T4; BioLegend), anti-CD8 Alexa700 (clone RPA-T8; BioLegend), anti-TCR γ/δ APC (clone B1; BioLegend), anti-CD161 BV421 (clone HP-3G10; BioLegend), anti-Vα7.2 PE-Cy7 (clone 3C10; BioLegend), anti-CD69 BV605 (clone FN50; BioLegend), anti-Ki67 PE (clone Ki67; BioLegend), anti-CD38 PE-CF594 (clone Ki67; BD Biosciences) and anti-HLA-DR APC-Fire750 (clone L243; BioLegend). A dump channel in FITC included the following antibodies: anti-CD14 (clone TuK4, LifeTechnologies), anti-CD19 (clone HIB19, BioLegend), anti-CD15 (clone HI98, BioLegend) and anti-CD16 (clone 3G8, BioLegend).

### Unsupervised analysis of flow cytometry data

Two dimensionality reduction methods based on a neighbouring graph approach were implemented, t-Distributed Stochastic Neighbour Embedding (t-SNE) [70] and Uniform Manifold Approximation and Projection (UMAP) [71]. t-SNE algorithm was performed on the Cytofkit platform [72] using up to 5,000 cells from each sample. UMAP was run as a plugin on FlowJo (v.10.4.1) using 15 nearest neighbours, a minimum distance of 0.5 and Euclidean distance for selected parameters. Files with .fcs extension from related experimental conditions were concatenated before UMAP analysis.

### Co-culture of *Salmonella* infected monocyte-derived dendritic cells and purified T cells

Monocyte-derived dendritic cells (MoDCs) were obtained from PBMC by enrichment of CD14^+^ monocytes using magnetic beads (Miltenyi). Differentiation was achieved with recombinant human GM-CSF (40 ng/mL) and human IL-4 (40ng/mL, both from PeproTech). After 5 days, MoDCs were infected with violet-labelled (CellTracker™, Life Technologies) *Salmonella* strains, either STM-D23580 or STM-LT2 at an MOI of 10, as reported elsewhere [42].

At six hours post-infection, *Salmonella*-containing MoDCs were FACS sorted as single cells. Sorted MoDCs were co-cultured with magnetically enriched (Miltenyi) CD3^+^ T cells obtained from the same donor, at a ratio of 1 MoDC per 6 T cells. Following 12 hours incubation in the presence of brefeldin A, T cells were harvested and stained for intracellular cytokines as described above.

### Co-culture of MoDCs and expanded MAIT cells

Human MoDCs were obtained as described above. Human MAIT cells were isolated by sorting CD2 MACS-enriched (Miltenyi) leukocytes with CD161 and Vα7.2 antibodies (BioLegend). MAIT cells were grown for around 6 weeks in complete RPMI media supplemented with IL-2, as described elsewhere [69].

40,000 MoDCs and 20,000 MAIT cells (2:1 ratio) were seeded in 96 well plates flat bottom and were infected at MOI of 3.5. After 80 minutes, 100 μg/mL gentamicin was added and supernatants were harvested following 26 hours incubation. IL-12 p70 was measured by ELISA (R&D systems) in triplicates and following manufacturer’s instructions.

### MR1 over-expressing cell line

THP-1 cells were transduced with an MR1 encoding lentivirus [34]. MR1 overexpressing cells were seeded in 96 well plates flat bottom and incubated overnight in the presence of 50 μL of supernatants from bacterial cultures at late exponential phase, or in the presence of 5-A-RU as positive control. Cells were harvested, washed and stained for surface expression of MR1 (clone 26.5; BioLegend) by flow cytometry. Expression of MHC-I (clone G46-2.6, BD Biosciences) was also monitored as unrelated control.

### Transcriptomic and proteomic analyses of riboflavin enzymes

RNA-seq and proteomic data for genes involved in the riboflavin biosynthetic pathway were extracted from a recent work [43]. Briefly, a differential expression comparative analysis between strains STM-D23580 and STM-4/74 was performed at the transcriptomic level in five *in vitro* infection-relevant conditions: ESP (early stationary phase), anaerobic growth, NonSPI2 (SPI2-non-inducing PCN), InSPI2 (SPI2-inducing PCN) and inside murine RAW264.7 macrophages (ATCC TIB-71). Specific details about growing bacteria in these conditions had been previously described [40], [73]. For a comparative proteomic analysis, bacteria were grown to ESP in the LB rich medium.

The RNA-seq-based comparative approach between STM-D23580 and STM-4/74 was based on Voom/Limma analysis from three different biological replicates for each strain. A detailed pipeline for the analysis can be found in Canals *et.al.* [43].

Proteomic data for strains STM-D23580 and STM-4/74 were obtained using an LC-MS/MS (Q Exactive Orbitrap, 4h reversed phase C18 gradient) platform. Samples included six biological replicates for each strain. Label-free quantification and differential expression analyses between the two strains were performed using the Progenesis QI software (Nonlinear Dynamics) [43].

### Measurement of riboflavin, FMN and FAD in supernatants and pellets of bacterial cultures

Bacterial pellets and supernatants from early stationary phase cultures (OD_600_=∼2), were prepared in triplicate and frozen at −80 °C. For extraction of cellular flavins, pellets were resuspended in 100 μL of 100 mM ammonium formate, 100 mM formic acid, 25% (v/v) methanol and heated at 80 °C for 10 min. Insoluble material was removed by centrifugation. For analysis 5 μL of this material or the culture supernatants was separated by HPLC on a Dionex UPLC system with a Kinetex C18 column (Phenomenex; 1.7 um, 150 × 2.1 mm). Separation was achieved at 45 °C and 0.2 mL/min isocractically using 20 mM potassium phosphate buffer (pH 2.5) with 25% methanol (v/v) over 8 min followed by a 1 min wash step in 100% methanol. Flavins were detected with fluorescence (450 nm excitation, 520 nm emission) and peaks were quantified by comparison to known standards. For normalisation the total pmol of flavin for each culture was divided by the measured OD600 of the cultures to give a final pmol/OD_600_ value.

### Statistical analysis

Statistical analyses were performed using GraphPad-Prism7 (GraphPad Software; San Diego, United States). Differences among groups were determined by paired one-way or two-way ANOVA, as appropriate. Post-hoc corrections were applied, Dunnett’s for comparisons to a control data set and Bonferroni’s, Tukey’s or Sidak’s for comparisons of selected pairs tests, as appropriate. A *p*-value <0.05 was considered statistically significant (\**p* <0·05, \*\**p* <0·01, \*\*\**p* <0·001 and \*\*\*\**p* <0·0001).

## ACKNOWLEDGMENTS

This work was supported by an NIHR Research Professorship and a Wellcome Trust Investigator Award (A.S.), the UK Medical Research Council through the MRC Human Immunology Unit (G.N., M.S and A.S.) and Celgene (A.A.). Part of this work was supported by a Wellcome Trust Senior Investigator award (to JCDH) (Grant 106914/Z/15/Z). PM was supported by a BBSRC David Phillips Fellowship (Grant BB/S010122/1). We acknowledge support of the Oxford NIHR Biomedical Research Centre. The views expressed are those of the authors and not necessarily those of the NHS, the NIHR or the Department of Health. We thank Dr. Ted Hansen for the gift of the anti-MR1 26.5 Ab. We acknowledge the contribution of Paul Sopp and Craig Waugh in the flow cytometry facility at the Weatherall Institute of Molecular Medicine for cell sorting experiments. We acknowledge the assistance of Priyanka Patel, Joyce Macheso, Anstead Kankwatira, and the Malawi-Liverpool-Wellcome Trust clinical team.

## AUTHOR CONTRIBUTIONS

Conceptualization L.P-L. M.S. and A.S.; Writing-Review & Editing, L.P-L., M.S., R.C., J.C.D.H., G.N., M.A.G. and A.S.; Methodology, L.P-L., M.S., R.C., A.A., P.M. and G.N.; Investigation, L.P-L., A.A., R.C., P.M., X.Z., N.J. and I.K.; Visualization, L.P-L., M.S., R.C. and P.M; Resources J.C.D.H., S.V.O., N.V., G.S.B., M.A.G. and T.N.; Supervision and funding, M.S and A.S.

