## Supplemental material for "African *Salmonella* Typhimurium sequence type 313 lineage 2 evades MAIT cell recognition by overexpressing RibB"

Preciado-Llanes *et al*.

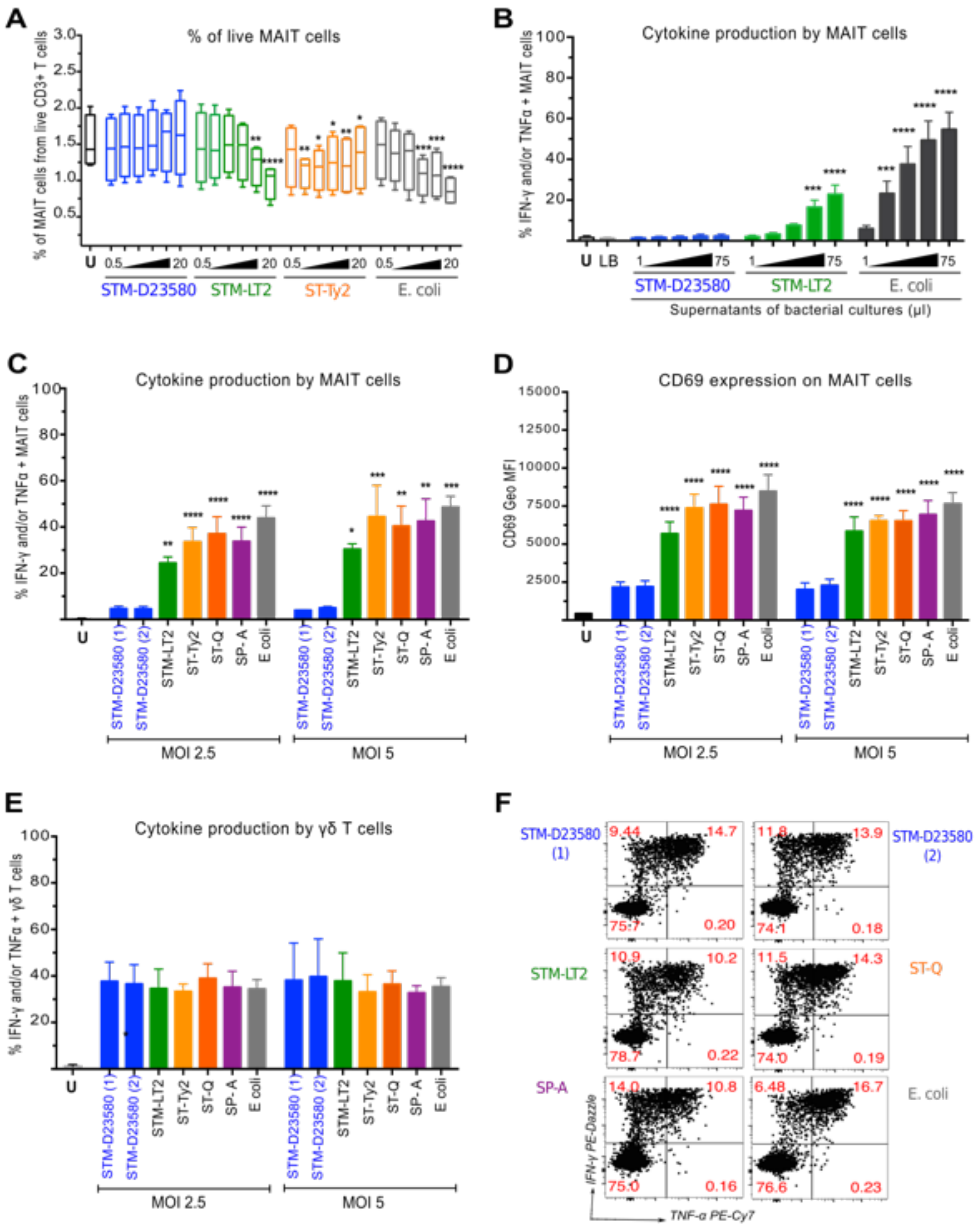


**Figure S1. STM-D23580 escapes from MAIT cell recognition without affecting cell viability and without reducing γδ T cell activation. (A)** PBMC were left unstimulated (U) or were infected with a variety of *Salmonella* strains at increasing MOI, from 0.5 to 20 bacteria per cell. *E coli* was included as positive control. Frequencies of MAIT cells from gated live $CD3^+$ T cells are plotted. Data represented as box-and-whisker plot, two-way ANOVA + Dunnet's (vs. STM-D23580), n=4. **(B)** Percentage of TNF-α and/or IFN-γ producing MAIT cells when stimulated with increasing amounts of bacterial culture supernatants, from 1 to 75 µL (final volume in well 250 µL). Data represented as mean ± SEM, two-way ANOVA + Dunnet's (vs. STM-D23580), n=4. **(C)** Percentage of TNF-α and/or IFN-γ producing MAIT cells, treated with bacterial strains at MOI of 2.5 and 5. Two STM-D23580 stocks were tested and are shown as (1) and (2). Data represented as mean ± SEM, one-way ANOVA + Dunnet's (vs. STM-D23580), n=5 for MOI 2.5 and n=3 for MOI 5. **(D)** Levels of CD69 expression on MAIT cells treated as in (C). Data represented as geometric mean ± SEM, one-way ANOVA + Dunnet's (vs. STM-D23580), n=5 for MOI 2.5 and n=3 for MOI 5. **(E)** Percentage of TNF-α and/or IFN-γ producing γδ T cells, treated with bacterial strains at MOI of 2.5 and 5. Data represented as mean ± SEM, one-way ANOVA + Dunnet's (vs. STM-D23580), n=5 for MOI 2.5 and n=3 for MOI 5. **(F)** Representative example of cytokine production by stimulated γδ T cells treated as in (E).

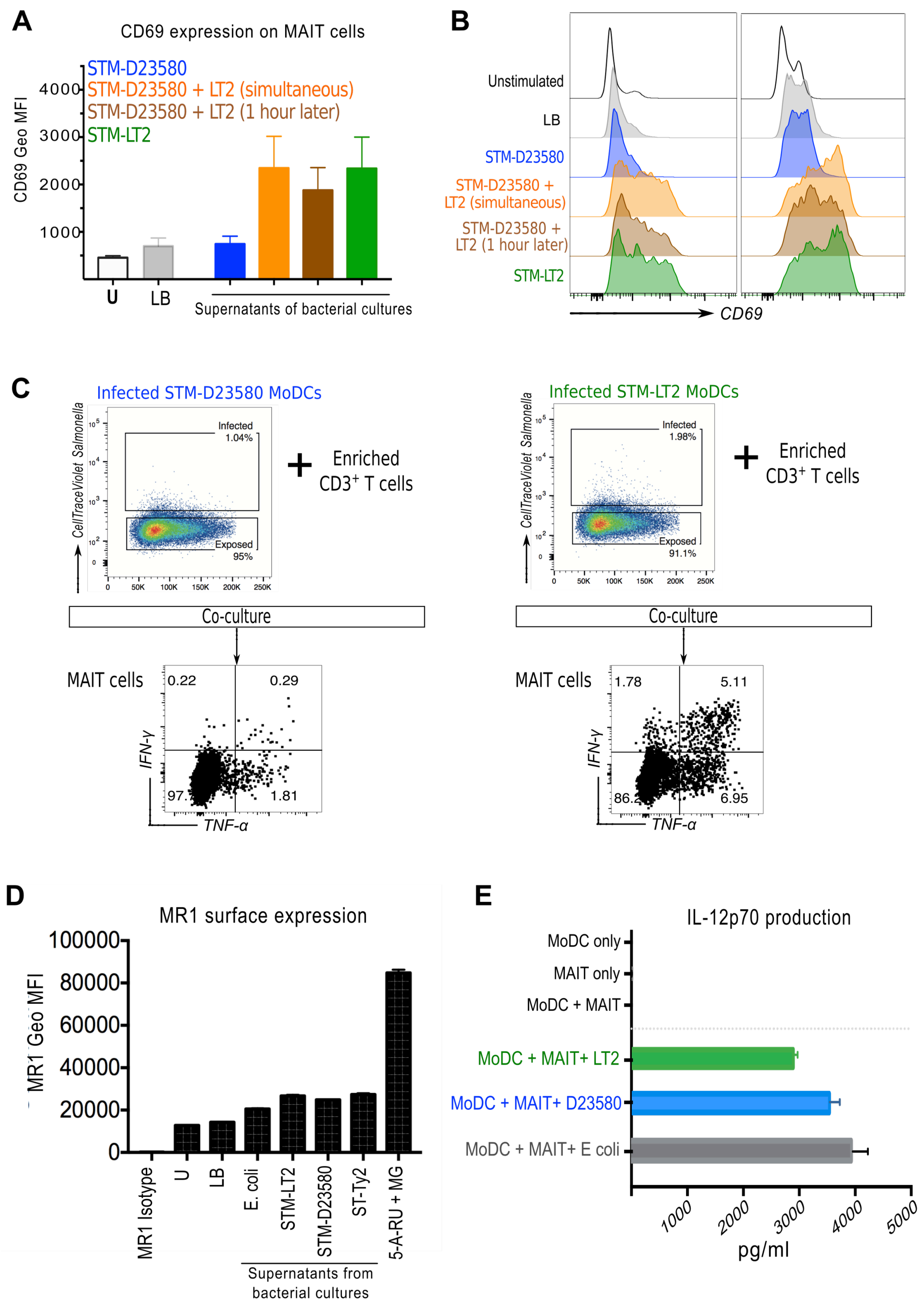


**Figure S2. STM-D23580 does not interfere with MR1-dependent presentation or with IL-12p70p production. (A)** CD69 expression on MAIT cells upon PBMC stimulation for 6 hours with 60 μL of STM-D23580 culture supernatant alone (blue bar) or in combination with 60 μL of STM-LT2 culture supernatant, either simultaneously (orange bar) or 1 hour apart (brown bar). LB medium (grey bar) and culture supernatant from STM-LT2 alone (green bar) were used as negative and positive controls, respectively. Data represented as geometric mean ± SEM, n=2. **(B)** Representative histograms from one volunteer treated as in (A) are shown. **(C)** Human monocyte-derived dendritic cells (MoDCs) were infected with violet-labelled (CellTracker™, Life Technologies) STM-D23580 or STM-LT2 at MOI of 10. At six hours post-infection, *Salmonella*-containing MoDCs (infected gate) were FACS sorted and co-cultured with autologous enriched $CD3^+$ T cells. Representative flow cytometry plots from one donor showing the percentage of TNF-$\alpha$ and/or IFN-$\gamma$ producing MAIT cells. **(D)** MR1 overexpressing cells were incubated overnight in the presence of 50 µL of supernatants from bacterial cultures as indicated. 5-A-RU + MG and LB medium were included as positive and negative controls, respectively. Surface expression of MR1 was assessed by flow cytometry. Data represents geometric mean ± SEM, 1 biological replicate in duplicates. **(E)** Human MoDCs were co-cultured with non-autologous expanded human MAIT cells and infected with STM-D23580, STM-LT2 or *E. coli* at MOI of 3.5. Supernatants were harvested following 26 hours incubation and IL-12p70 was measured by ELISA, 1 biological replicate in triplicates.

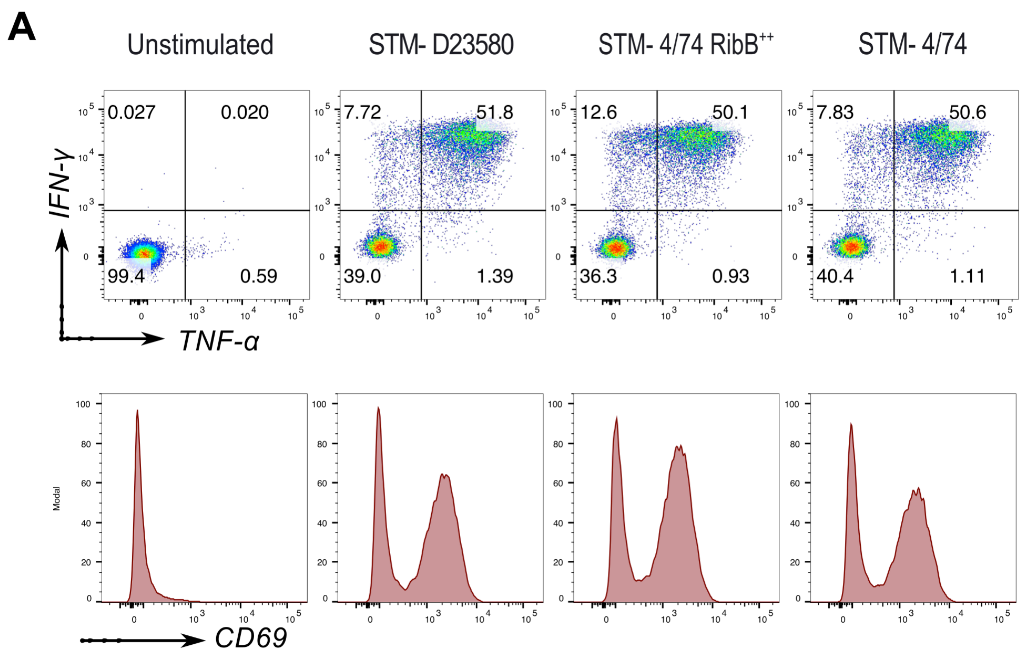


**Figure S3. γδ T cell responses to STM-4/74 RibB$^{++}$ are not different from STM-4/74 wild type.**

PBMC were infected at MOI of 2 with STM-D23580, STM-4/74 or STM-4/74 RibB$^{++}$. **(A)** Representative dot plots showing the percentage of TNF-α and/or IFN-γ producing γδ$^+$ T cells. **(B)** Representative histograms for CD69 expression on γδ$^+$ T cells from the same donor as in (A).

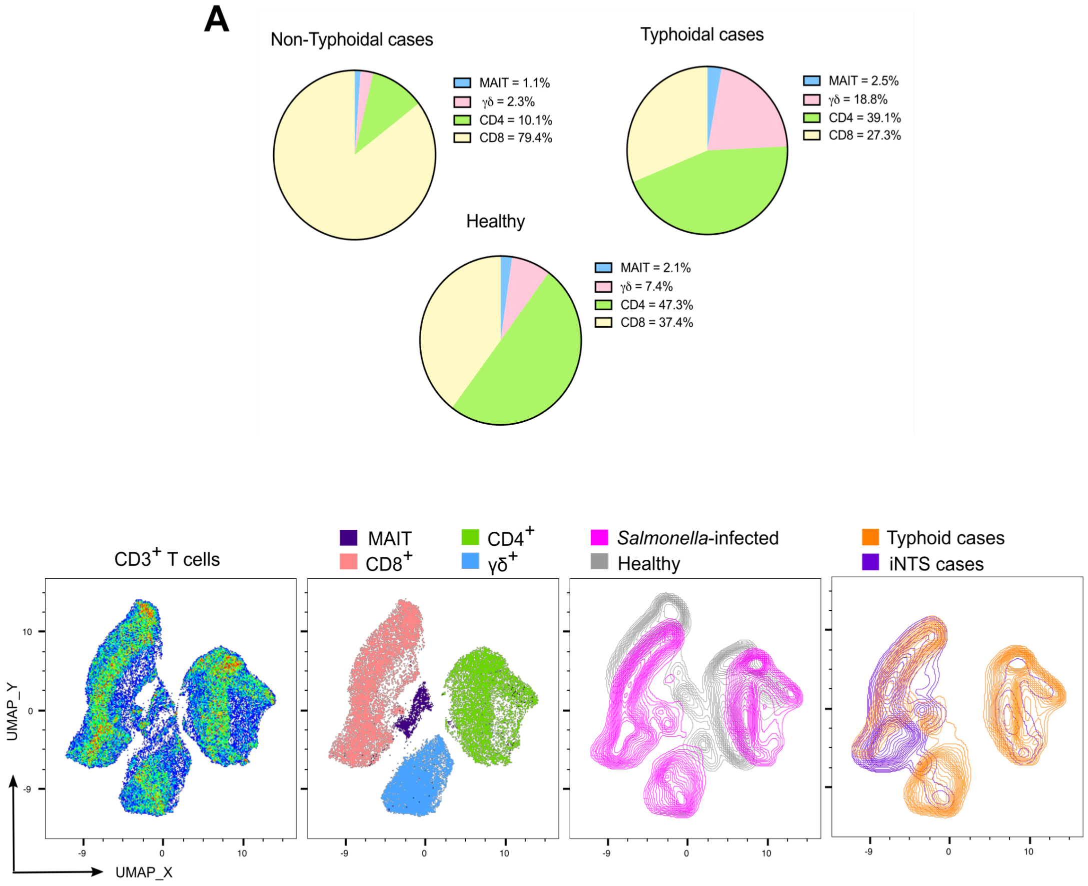


**Figure S4. Characterisation of MAIT cells in a cohort of patients with Salmonellosis.**

Whole blood aliquots, from typhoidal (n=5) and nontyphoidal cases (n=3), as well as healthy individuals living in Malawi (n=4) were stained for T cell markers and analysed by flow cytometry. **(A)** Proportion of $CD4^+$, $CD8^+$, MAIT cells ($CD161^+$ $V\alpha7.2^+$) and $TCR\gamma\delta^+$ T cells are reported as pie charts using average percentages by study group. **(B)** UMAP analysis was performed on concatenated manually gated $CD3^+$ T cells (75,000 cells in total) from cases and healthy controls. The different T cell subsets are shown by colours on the dot plot. The distribution of healthy, iNTS and typhoid cases is shown as overlay contour plots.

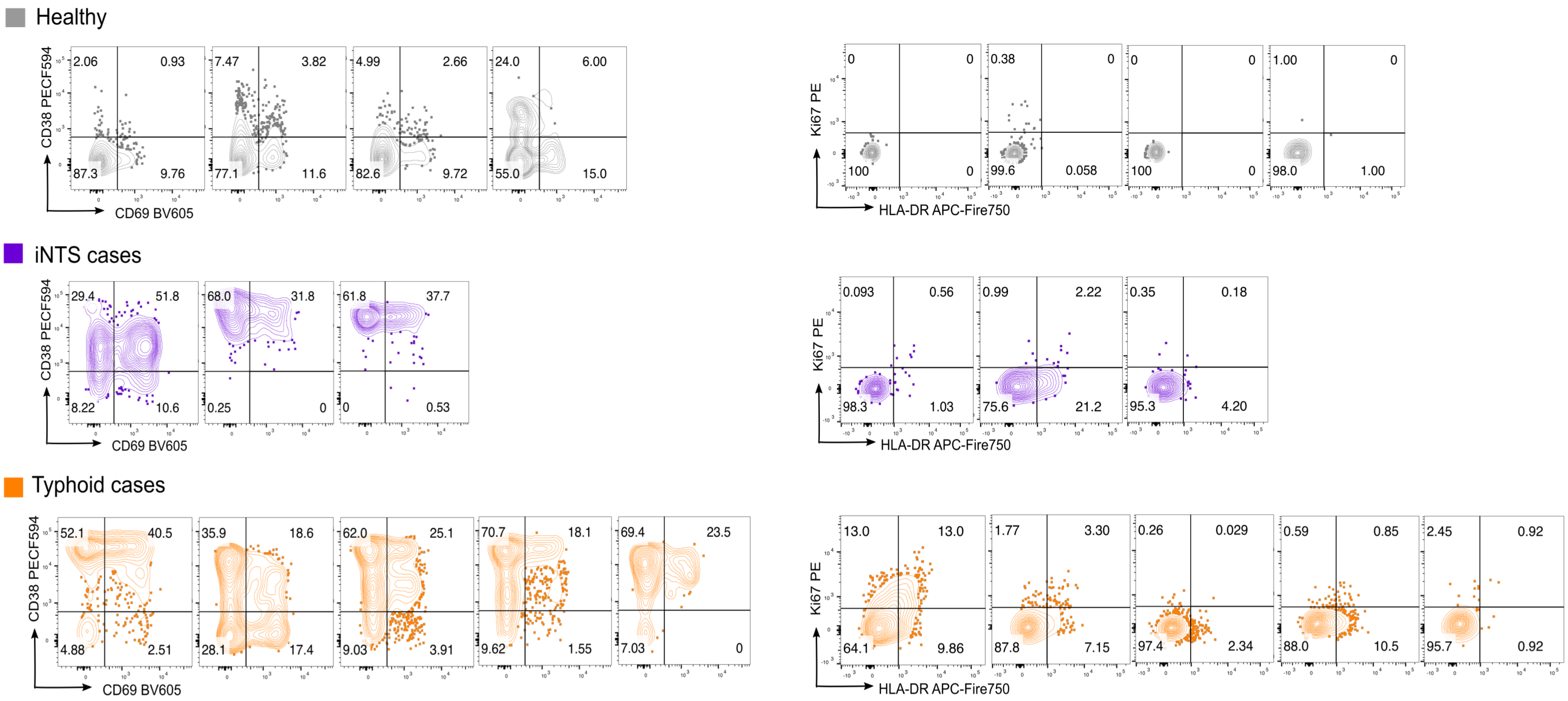


**Figure S5. Activation phenotype of MAIT cells in a cohort of patients with Salmonellosis.**

MAIT cells gated from cases and healthy control samples as from Figure S4. Flow cytometry dot plots show the percentage of MAIT cells expressing a combination of activation markers: CD38, HLA-DR, CD69 and Ki67.

**Supplementary Table 1.**

| Resource | Source | Identifier |
|---|---|---|
| **Bacterial Strains** | | |
| *Salmonella* Typhimurium 4/74 (ST19) | (Rankin and Taylor, 1966) | JH3676 |
| *Salmonella* Typhimurium LT2 (ST19) | (McClelland et al., 2001) | ATCC 700220 |
| *Salmonella* Typhimurium D23580 (ST313 lineage 2) | (Kingsley et al., 2009) | JH3621 |
| *Salmonella* Typhimurium D25248 (ST313 lineage 1) | (Kingsley et al., 2009) | IC24T |
| *Salmonella* Typhimurium U60 (ST313 lineage 2) | (Ashton et al., 2017) | IC39S |
| *Salmonella* Typhimurium U5 (UK-ST313 strain) | (Ashton et al., 2017) | IC25I |
| *Salmonella* Typhimurium U2 (UK-ST313 strain) | (Ashton et al., 2017) | IC25F |
| *Salmonella* Typhimurium D37712 (ST313 lineage 2) | (Msefula et al., 2012) | IC24O |
| *Salmonella* Typhi Ty2 | (Deng et al., 2003) | ATCC 700931 |
| *Salmonella* Typhi Quailes | (Waddington et al., 2014) | NA |
| *Salmonella* Paratyphi NVGH308 | (Dobinson et al., 2017) | NA |
| *E. coli* DH5$\alpha$ | ThermoFisher | NA |
| *E. coli* TOP10 (*mcrA D(mrr-hsdRMS-mcrBC)* $\phi$8 0*lac*ZDM15 *ΔlacX74 deoR recA1 araD139* Δ*(ara-leu)7697galU galK rpsL endA1 nupG*) | Invitrogen | JH4317 |
| *E. coli* TOP10 pP$_L$-*ribB* | This study | JH4318 |
| *S.* Typhimurium 4/74 pP$_L$-*ribB* | This study | JH4319 |

**Supplementary Table 2.**

| **Plasmids** | **Source** | **Identifier** |
|---|---|---|
| ColE1 control plasmid, based on pZE12-luc, $P_{LlacO}$ promoter transcribes a ~50 nt nonsense transcript (*rrnB* terminator) | (Sittka et al., 2007) | pJV300 |
| pJV300 ($pP_L$) plasmid carrying the coding region of the *ribB* gene | This study | $pP_L$-*ribB* |
| **Oligonucleotides** | | |
| GTGAGCGGATAACAAGATACTGAGCACCTGGTAACCA TAATATTAATGAGG | This study | *ribB*_FW |
| GCCTTTCGTTTTATTTGATGCCTCTAGAATCAGCTGGC TTTGCGCTCATGCG | This study | *ribB*_RV |
| GTGCCACCTGACGTCTAAGA | This study | pPL_Seq_FW |
| ATACCGCTCGCCGCAGCCG | This study | pPL_Seq_RV |
